# BugBuilder - An Automated Microbial Genome Assembly and Analysis Pipeline

**DOI:** 10.1101/148783

**Authors:** Abbott J.C.

**Affiliations:** Bioinformatics Data Science Group, Department of Surgery and Cancer, Imperial College London, London, UK, SW7 2AZ

## Abstract

**Summary:** BugBuilder is a framework for hands-free assembly and annotation of microbial genomes. It produces outputs suitable either for database submission or downstream finishing processes. It is configurable to work with most command-line assembly and scaffolding tools which are selectable at run-time, and supports all common sequence types used in microbial genome assembly.

**Availability and Implementation:** BugBuilder is implemented in Perl and is available under the Artistic License from http://www.imperial.ac.uk/bioinformatics-data-science-group/resources/software/bugbuilder, A virtual machine image is available pre-configured with the relevant freely-redistributable dependencies.

## 1 Introduction

The rapid increase in sequencing throughput in recent years has moved the bottleneck in the process of sequencing microbial genomes from sequence generation to the analysis. The cost of sequence analysis now typically outweigh those of sequencing, while the use of multiplexing combined with the scale of modern sequencing instruments typically results in large batches of sequences being produced simultaneously. Lab-based researchers often do not have the background knowledge necessary to carry out the required analysis themselves.

BugBuilder provides a portable framework for carrying out assembly and analysis of microbial genome sequences, taking sequence reads as inputs and producing submission-ready annotated genome assemblies, with ease-of-use prioritised to allow non-expert users to obtain acceptable results. The software selects appropriate tools and parameters based upon the type of sequence data provided. It is also suitable for more advanced applications, being designed to be readily deployed in a cluster environment while permitting manual run-time selection of tools and parameters, and can be readily customised to include any Linux command-line based assembly and scaffolding tools. the outputs of most major sequencing platforms are supported, including long-read and hybrid assemblies, and can either provide submission-ready outputs, or augment these with indications of potential sites of misassembly to assist with downstream finishing work.

## 2 Implementation

BugBuilder is implemented in Perl, utilising a YAML format configuration file. A BugBuilder job carries out it’s work in a temporary directory, within which a separate directory is created for each component of the pipeline. A series of symbolic links are maintained within the working directory which link to, for example, the most recent version of generated contig sequences, facilitating flexible workflows through consistent file naming

In order to prevent the system being tied to particular software tools or sequencing platforms, multiple assemblers and scaffolders are defined with the central configuration. Any package which can be run on the command-line can be integrated into BugBuilder, either directly where the tools input and output requirements are appropriate for BugBuilder, or through the use of a script wrapper where more complex requirements exist. Default arguments to the tools are also defined in the configuration, which can be overridden at run time through command-line arguments.

## 3 Workflow

The BugBuilder workflow is shown in Figure 1. The required inputs are fastq files obtained from a sequencing experiment, where a single fastq file is required from a fragment library, or a pair of non-interleaved fastq files in the case of a mate-pair library. Sequence from long-read platforms (i.e. PacBio, MinION) can either be used for standalone assembly or combined with short-read sequences for a hybrid assembly. A fasta-formatted reference genome sequence can optionally be provided which may be used for scaffolding and ordering scaffolds if desired. Assemblers and scaffolders are categorised according to applicable sequencing platforms, so the user can either define the sequencing platform used and allow BugBuilder to select the most appropriate tools based on the sequence type, or manually select the desired programs from the applicable tools defined for the sequence category.

**Figure 1:**
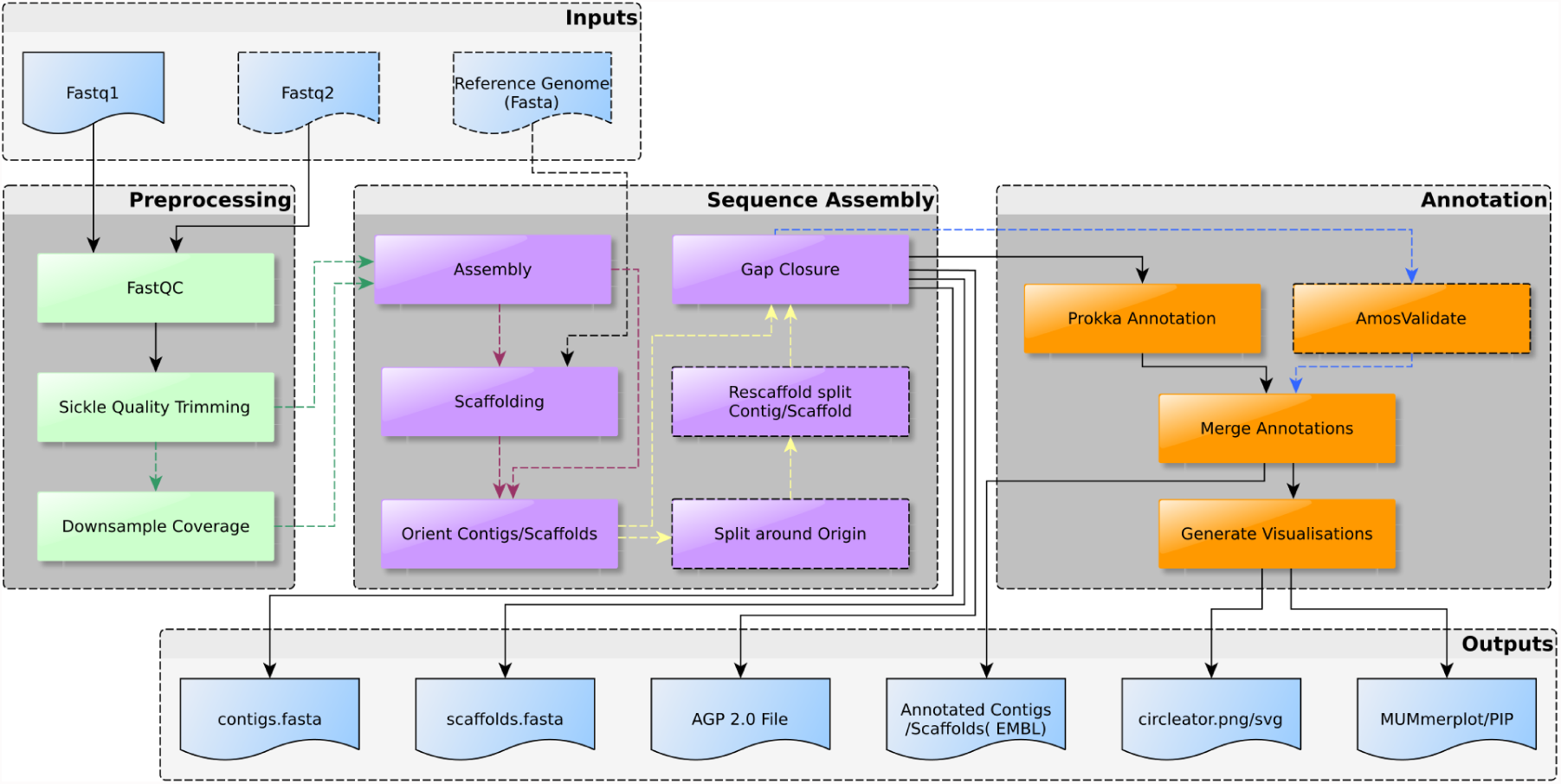
The BugBuilder Workflow. Alternate paths through the workflow are indicated by arrow colour, with dotted arrows and borders indicating conditional stages and routes.

### 3.1 Sequence Assessment and Preprocessing

Prior to assembly, a number of preprocessing stages are carried out. The processes executed vary according to sequence type, and individual stages can be skipped if desired by the user. Sequence reads are inspected to determine the sequence characteristics, sequence quality is assessed using FastQC (http://www.bioinformatics.babraham.ac.uk/projects/fastqc/), followed by sequence trimming using Sickle (https://github.com/najoshi/sickle) to remove low quality bases for assemblers which do not carry out trimming directly. High coverages in *de Bruijn* genome assemblers can lead to a decrease in assembly quality, consequently reads are downsampled to either 100x or a user-defined threshold for such assemblers.

### 3.2 Contig Assembly

The default BugBuilder configuration includes the ABySS (Simpson *et al.*, 2009), SPAdes (Bankevich *et al.*, 2012) and Celera WGS (CABOG and PBcR) (Miller *et al.*, 2008, Berlin *et al.*, 2015) assemblers, providing support for sequence obtained from Illumina short-read platforms (i.e. GAII, HiSeq, MiSeq), through to Roche 454, PacBio and MinION instruments. The assembly stage produces, at minimum, a fasta file of contig sequences, and optionally a fasta file of scaffolded sequences where scaffolding is supported directly by the assembler.

### 3.3 Scaffolding

BugBuilder considers two classes of scaffolder; those which utilise read-pair associations between contigs, and those which carry out alignment to a reference sequence. Reference-free methods are less susceptible to biasing the outputs to the organisation of the reference sequence, however typically produce considerably less contiguous results, consequently the preferred algorithm is highly dependent upon similarity to the reference genome and user preference. Predefined configurations for scaffolding with SSPACE (Boetzer *et al.*, 2011), SIS (Dias *et al.*, 2012) and the Mauve Contig Mover (Rissman *et al.*, 2009) are provided with the software.

Scaffolds can frequently be further improved by ordering and orientating them against the reference genome sequence, especially if a reference-free scaffolding algorithm was used. The origin of replication will typically occur within a contig sequence, although by convention this is used as the start of the genome sequence. The location of the origin is determined by alignment against the reference (if provided), and the assembly orientated around this locus. Remaining gaps between contigs can then be processed with GapFiller (Boetzer and Piravano, 2012), which carries out incremental alignment of sequence reads around contig ends to extend contigs and close gaps.

### 3.4 Annotation, Visualisation and Outputs

Annotation of assembled sequences is carried out using Prokka (Seemann, 2014), which combines various feature predictions algorithms and similarity search methods to produce an annotation including coding genes, rRNAs and tRNAs. BugBuilder also allows execution of a downstream validation process (amosvalidate - Phillippy *et al.*, 2008) to help identify potential misassemblies by evaluating read pair distribution to identify compressed or extended read-pairs and inversions etc., and read depth to identify collapsed or expanded repeats.

The main output of the pipeline is an EMBL-format record containing the annotated scaffold or contig sequences which is appropriate for submission to the ENA. Visualisation and interpretation of results are aided by the production of a graphical genome map using Circleator(Crabtree *et al.*, 2014), along with similarity plots and percentage-identity plots generated with MUMmerplot (Kurtz *et LA.*, 2004). A full list of outputs generated is provided in table 1.

**Table 1:** Output files produced by BugBuilder. Outputs vary according to executed tools as indicated in ‘When produced’ column as follows: ‘Always’ - output is always produced; ‘If scaffolded’ - produced if the assembly is scaffolded, either by the assembler or a scaffolder being run; ‘Unless scaffolded’ - produced if the assembly is not scaffolded; ‘With reference’ - output only produced if a reference sequence is provided.

| Filename | Type | When produced | Description |
| --- | --- | --- | --- |
| read[12]_fastqc.html | HTML | Always | Output of FastQC quality assessment |
| contigs.embl/gb | EMBL/GenBank | Unless scaffolded | Annotated contig sequence in submission-ready format. Amosvalidate results (if present) will be represented as misc_features. |
| contigs.fasta | Fasta | Always | Contig sequences in fasta format |
| unplaced_contigs.fasta | Fasta | If scaffolded | Contig sequences which could not be incorporated during reference-based scaffolding |
| scaffolds.embl/gb | EMBL/GenBank | If scaffolded | Annotated scaffold sequences in submission-ready format. Amosvalidate results (if present) will be represented as misc_features. |
| scaffolds.fasta | Fasta | If scaffolded | Scaffold sequences in fasta format |
| scaffolds.agp | AGP 2.0 | If scaffolded | Details of scaffold construction |
| comparison_vs_[accession].png | PNG image | With reference | MUMmerplot output of nucmer alignment between scaffolds and reference, indicating structural differences between reference and sample |
| comparison_vs_[accession]_pip.png | PNG image | With reference | MUMmerplot percentage identity plot, indicating sequence similarity between reference and sample |
| comparison_vs_[accession].blastout | NCBI Blast output | With reference | Alignment between sample and reference with NCBI Blast, formatted for visualisation with ACT |
| circleator.png | PNG image | Always | Circleator genome map |
| circleator.svg | SVG image | Always | Circleator genome map |

### 3.5 Example Assemblies

Sequence data for E. coli K-12 strains generated from a range of platforms was obtained from the ENA database and used to validate the pipeline outputs (see table 2). Each was assembled using the default assembler, both with and withough reference-based scaffolding using SIS, with just the platform, reference sequence (U00096.3) and choice of scaffolder (where required) being provided as command-line arguments. Assemblies were run using 8 threads on AMD Opteron 6282 SE 2.6Ghz CPUs, with 8Gb RAM available per-core.

**Table 2:** E.coli K-12 libraries used for assemblies.

| Platform | Organism | ENA Accession | Read Type | Insert Size | Coverage |
| --- | --- | --- | --- | --- | --- |
| Illumina GA | E. coli K-12 substr. MG1655 | PRJNA30551 | 37 bp paired | 486 bp | 112x |
| Illumina HiSeq 2000 | E. coli K-12 substr. MG1655 | PRJEB4687 | 102 bp paired | 240bp | 810x |
| Illumina MiSeq | E. coli K-12 | PRJEB8559 | 128 bp paired | 4914 bp | 49x |
| Roche 454 FLX | E. coli K-12 substr. MG1655 | PRJNA40075 | Unpaired, mean 228 bp | - | 29x |
| PacBio RSII | E. coli K-12 substr. MG1655 | PRJNA237120 | Unpaired long reads, mean 1954 +/- 2621 bp | - | ~139x |

BugBuilder has to date been successfully used on projects involving a range of species including Streptococcus pyogenes (e.g. Turner *et al.*, 2015), Escherichia coli, Gluconacetobacter hansenii (e.g. Florea *et al.*, 2016) and Klebsiella pneumoniae.

## 4 Assembly Results

The outputs of the test assemblies are shown in table 3 and figures 2-5. As would be expected the main factor determining the contiguity of assembly is the sequencing technology employed and the subsequent choice of assembly and scaffolding algorithm. The configurable nature of the pipeline readily permits the addition of new algorithms with improved capabilities or technology support.

**Table 3:** E. coli K-12 assembly results.

| Platform | Assembler | Scaffolder | GapFilled | Assembly Size (bp) | Contigs (>200 bp) | Contig N50 (bp) | Scaffolds | Scaffold N50 (bp) | CDS |
| --- | --- | --- | --- | --- | --- | --- | --- | --- | --- |
| Illumina GA | SPAdes | - | No | 4537545 | 396 | 25549 | 166 | 65561 | 4253 |
|  | SPAdes | SIS | No | 4548329 | 416 | 23897 | 8 | 3672437 | 4245 |
| Illumina HiSeq 2000 | SPAdes | - | Yes | 4456532 | 114 | 108958 | 111 | 132988 | 4140 |
|  | SPAdes | SIS | Yes | 4495258 | 91 | 133146 | 4 | 3627322 | 4180 |
| Illumina MiSeq | SPAdes | - | Yes | 4561241 | 45 | 470241 | 44 | 565835 | 4274 |
|  | SPAdes | SIS | Yes | 4562536 | 36 | 470507 | 8 | 4558614 | 4272 |
| Roche 454 FLX | Celera WGS | - | No | 4555044 | 77 | 117927 | 76 | 117927 | 4245 |
|  | Celera WGS | SIS | No | 4562444 | 77 | 117929 | 2 | 2436314 | 4249 |
| PacBio RSII | PBcR | - | No | 4674981 | 2 | 3442086 | 1 | 4674981 | 4824 |
|  | PBcR | SIS | No | 4653988 | 2 | 3442086 | 1 | 4653988 | 4749 |
| Hybrid PacBio/MiSeq | SPAdes | - | No | 4560188 | 46 | 426752 | 45 | 565835 | 4276 |
|  | SPAdes | SIS | Yes | 4555687 | 34 | 669128 | 8 | 4551326 | 4275 |

**Figure 2:**
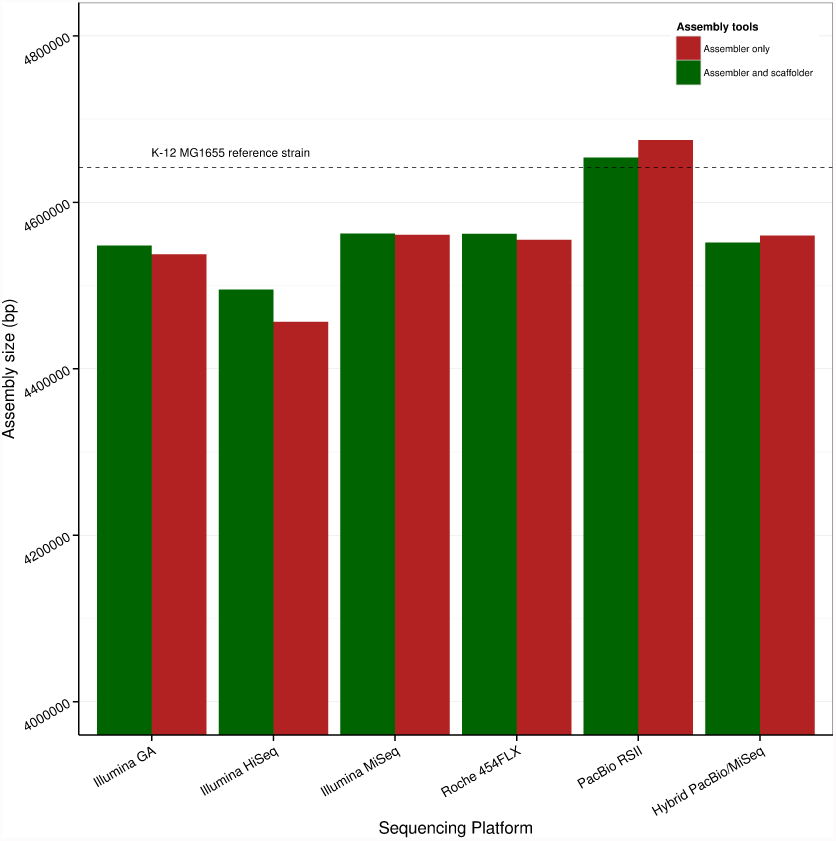
Total assembly sizes for E. coli K12 MG1655 example assemblies.

**Figure 3:**
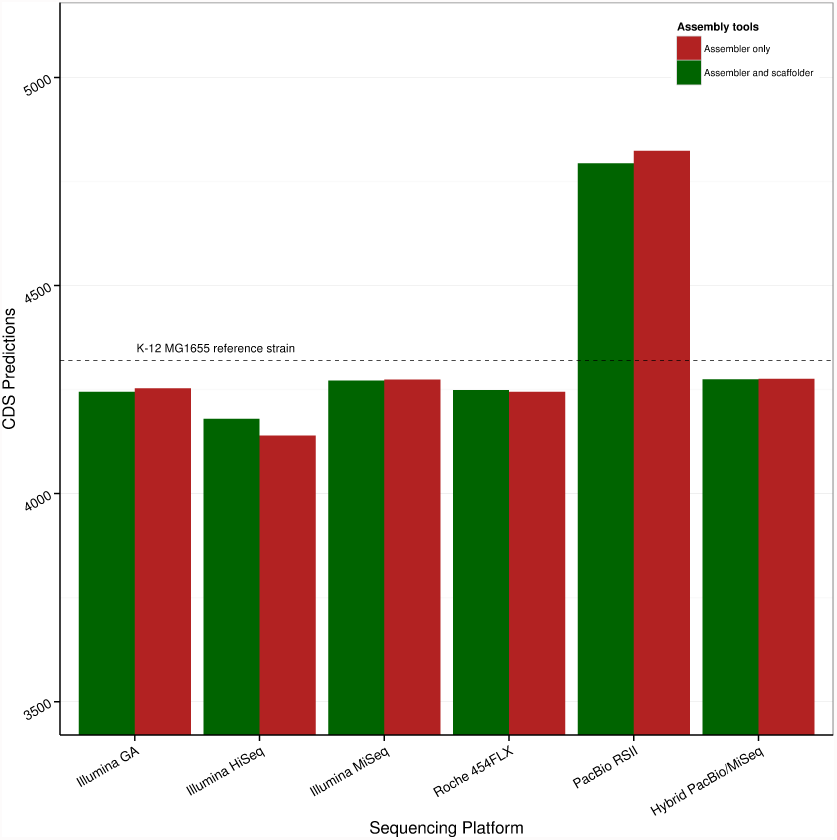
Number of CDS predictions from E. coli K12 MG1655 example assemblies. CDS predicitions were carried out on the scaffolding sequences resulting from each assembly using Prokka

**Figure 4:**
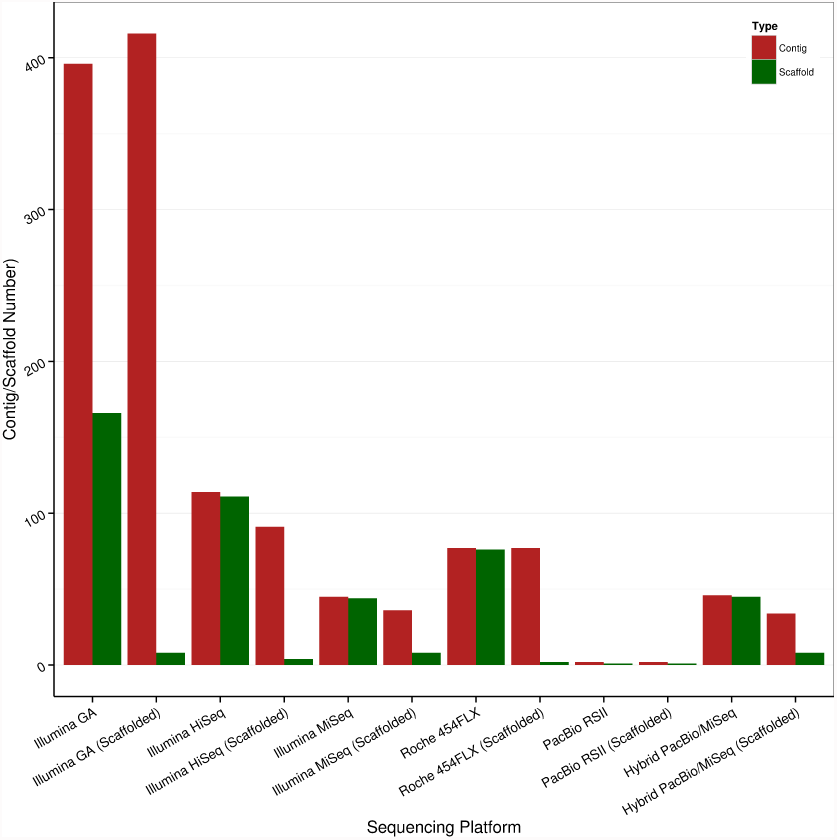
Numbers of contig and scaffold sequences resulting from E. coli K12 MG1655 example assemblies. Assembler and scaffolder selections used for each assembly are indicated in table 3.Statistics for scaffold sequences are either based upon scaffolds output by the assembler, or for the outputs of a downstream scaffolding process in assemblies labelled ‘Scaffolded’.

**Figure 5:**
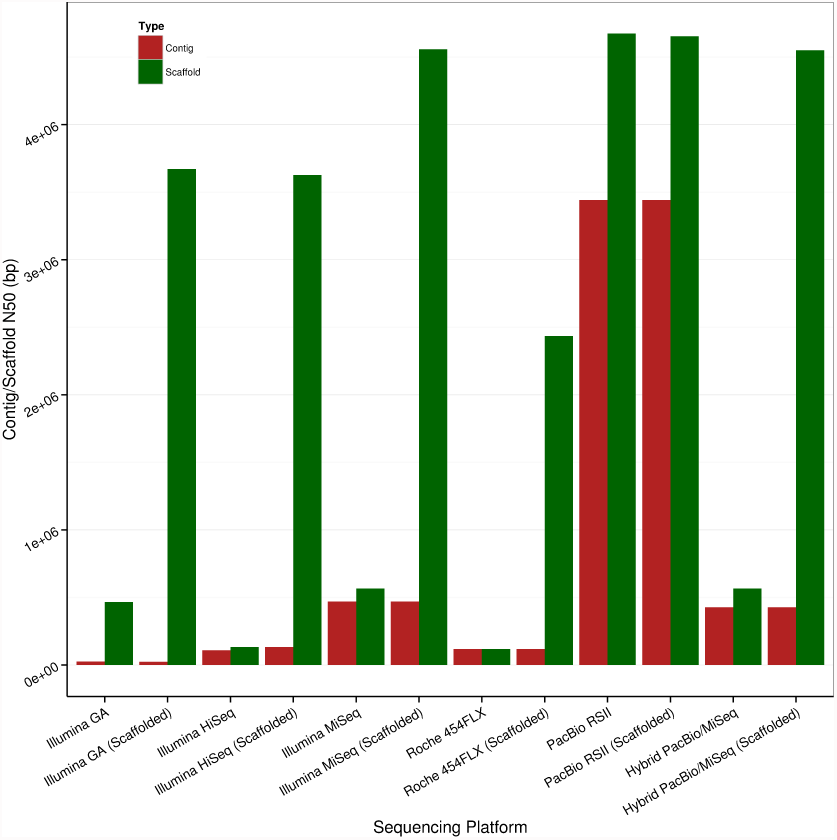
Contig and scaffold N50 values resulting from E. coli K12 MG1655 example assemblies. Assembler and scaffolder selections used for each assembly are indicated in table 3. Statistics for scaffold sequences are either based upon scaffolds output by the assembler, or for the outputs of a downstream scaffolding process in assemblies labelled ‘Scaffolded’.

